# LONG-TERM NEURODEVELOPMENTAL AND COGNITIVE OUTCOMES FOLLOWING PRENATAL INHIBITION OF DYRK1A BY LEUCETTINE L41 IN MOUSE MODELS OF DOWN SYNDROME

**DOI:** 10.64898/2026.05.13.724917

**Authors:** Arnaud Duchon, Claire Chevalier, Patrick Gizzi, Julien Dairou, Yann Herault

## Abstract

Down syndrome (DS), caused by trisomy of human chromosome 21, is characterized by intellectual disability and cognitive deficits, partly driven by the overexpression of Dual-specificity tyrosine-(Y)-phosphorylation Regulated Kinase 1A (DYRK1A). While postnatal DYRK1A inhibition has shown promise in improving cognition in DS models, its therapeutic potential during embryonic development, a critical window for neurogenesis, remains unexplored.

Here, we tested the hypothesis that prenatal inhibition of DYRK1A could mitigate long-term cognitive impairments in DS. We administered Leucettine L41, a potent and selective DYRK1A inhibitor, to pregnant dams carrying two DS mouse models: Ts65Dn and Dp(16)1Yey, both of which recapitulate trisomy of genes homologous to human chromosome 21, including *Dyrk1a*. Treatment was designed to suppress DYRK1A kinase activity during embryogenesis.

In adulthood, we evaluated the progeny for cognitive performance, gene expression profiles linked to DS phenotypes, and neuronal maturation markers. Prenatal L41 treatment produced lasting effects in both models, rescuing specific behavioral deficits and modulating the expression of DS-implicated genes, including the excitatory/inhibitory balance regulator GAD67. However, model-specific responses were observed: hyperactivity, working memory deficits, and GAD67-positive cell counts remained uncorrected in Ts65Dn mice, suggesting divergent molecular pathways underlying shared DS phenotypes.

This study demonstrates the therapeutic potential of prenatal DYRK1A inhibition for DS and provides novel insights into its role in neurodevelopmental trajectories and cognitive outcomes. Our findings underscore the importance of timing and genetic context in DS intervention strategies.

## INTRODUCTION

Down syndrome (DS) results from the presence of three chromosomes 21 in individuals and is the main genetic cause of intellectual disabilities (ID). It accounts for about 30% of the general population’s moderate to severe cases of ID (Antonarakis *et al*. 2004). Intellectual disability, present in 100% of individuals with DS, arises from developmental and functional alterations in the brain, leading to lifelong cognitive and adaptive challenges. To elucidate the neurobiological mechanisms underlying DS and to develop therapeutic strategies, mouse models carrying partial duplications of regions homologous to Hsa21 have been generated. These models recapitulate key neurological and cognitive phenotypes observed in humans with DS.

Among the most widely used models, the Ts65Dn line [Ts(17/16)65Dn] (Reeves *et al*. 1995) is the oldest and most studied. This model carries a supernumerary mini-chromosome, resulting in trisomy for 122 protein-coding genes located on mouse chromosome 16 (Mmu16), from *Mrpl39* to *Zbtb21* (Duchon *et al*. 2011; MuÑiz Moreno *et al*. 2020). While Ts65Dn mice exhibit many DS-associated phenotypes, a significant limitation is the presence of an additional trisomic region homologous to human chromosome 6, encompassing 45 non-DS-related genes. This extra trisomy exacerbates phenotypic severity, complicating the interpretation of results (Duchon *et al*. 2021a; Duchon *et al*. 2022). Nevertheless, studies in Ts65Dn mice have revealed critical defects in neurogenesis, particularly in the neocortical subventricular zone and the dentate gyrus during embryogenesis (Belichenko *et al*. 2004; Chakrabarti *et al*. 2007). These defects include reduced production of excitatory neurons, leading to an imbalance in the excitatory/inhibitory neuronal ratio (Chakrabarti *et al*. 2010). Additionally, neurogenesis and synaptogenesis are impaired in the hippocampus, with neuronal deficits stemming from proliferation abnormalities in precursor cells, from the neocortical subventricular zone and of the dentate gyrus, characterized by prolonged G2 and shortened M phases of the cell cycle (Chakrabarti *et al*. 2007; Contestabile *et al*. 2007).

Another key DS model, the Dp(16Lipi-Zbtb21)1Yey (noted here Dp1Yey) model (Li *et al*. 2007), carries a duplication of all genes in the Mmu16 region homologous to Hsa21, from *Lipi* to *Zbtb21* (Li *et al*. 2007). Unlike Ts65Dn, Dp1Yey does not harbor an extra chromosome, making it a more genetically precise model. This model also exhibits cognitive deficits and neurogenesis impairments similar to those observed in humans, though fewer neurodevelopmental alterations have been described (Goodliffe *et al*. 2016).

The phenotypes associated with DS are largely attributed to the overexpression of dosage-sensitive genes present in three copies (Antonarakis *et al*. 2004). Among these, the dual specificity tyrosine phosphorylation-regulated kinase 1A (DYRK1A) has gathered significant attention due to its pivotal role in brain development and function. Overexpression of *Dyrk1a* in DS models contributes to cognitive impairments reminiscent of those observed in individuals with DS (Duchon and Herault 2016). DYRK1A is a critical regulator of neurogenesis and neuronal maturation, influencing neuronal proliferation and differentiation (Hammerle *et al*. 2011; Soppa *et al*. 2014). Transient expression of *Dyrk1a* maintains neural precursors in a quiescent state, poised for differentiation (Hammerle *et al*. 2011). Furthermore, DYRK1A regulates the G1 phase of the fibroblast cell cycle, and its inhibition can reverse the prolonged G1 phase observed in DS mouse fibroblasts (Chen *et al*. 2013). Genetic normalization of *Dyrk1a* dosage in Ts65Dn mice has been shown to rescue certain DS-associated phenotypes(Garcia-Cerro *et al*. 2017).

Over the past decade, numerous DYRK1A inhibitors have been developed (Becker et al. 2014; Atas-Ozcan et al. 2021). Prenatal or antenatal administration of these inhibitors has been explored as a strategy to mitigate the adverse effects of DYRK1A overexpression during gestation. Early intervention during in-utero development aims to reduce the impact of DYRK1A overdosage on brain development, potentially improving cognitive outcomes in adulthood.

Pioneering studies demonstrated the efficacy of prenatal intervention using the natural DYRK1A inhibitor epigallocatechin gallate (EGCG). In the Tg(CEPHY152F7)12Hgc DS mouse model, which overexpresses human DYRK1A and three other genes, EGCG treatment initiated at the onset of pregnancy and continued into adulthood rescued major transgenic phenotypes (Guedj *et al*. 2009). Similar results were obtained in Dp1Yey mice, where prenatal EGCG treatment corrected working memory deficits (Delabar *et al*. 2024). These results were also confirmed by another study in the Tg(Dyrk1a)189N3Yah, overexpressing *Dyrk1a* alone (Guedj *et al*. 2009). Additionally, prenatal EGCG administration in Tg(Dyrk1a)189N3Yah mice, which overexpress *Dyrk1a* alone, and in Dp(16)1Yey mice, corrected the increased density of GAD67 neurons and hippocampal GAD67 overexpression (Souchet *et al*. 2019). In addition, when the treatment was stopped at weaning, the correction was maintained in the animal up to the age of 69 days, whereas only treating adult mice did not modulate those phenotypes (Souchet *et al*. 2019). ALGERNON, another DYRK1A competitive inhibitor, could also rescue some DS phenotypes after in utero treatment in the Ts1Cje mouse model, containing a smaller triplicated region homologous to human chromosome 21, going from *Sod1* to *Zbtb21* (Nakano-Kobayashi *et al*. 2017). The effects of postnatal inhibition of DYRK1A were previously investigated with the EGCG administrated to Ts65Dn mice from postnatal day P3 to P15 (Stagni *et al*. 2016). It allowed the restoration of the number of neural precursor cells in two major brain neurogenic regions in trisomic model. They also found that the deficit in the number of proliferative cells in the Ts65Dn disappeared with the EGCG prenatal treatment (Stagni *et al*. 2016). Unfortunately, the treatment failed to induce long-term effects in working and spatial memory. Conversely, a recent longitudinal study by Llambrich et al. (2024) found that chronic prenatal treatment with green tea extract-EGCG (Llambrich *et al*. 2024) exacerbated trisomic phenotypes in adult cognition, casting doubt on its therapeutic utility. The contradictory outcomes across studies may stem from several factors. First, the treatment window varies significantly, potentially influencing developmental trajectories. Second, the cognitive assessments employed differ in methodology and focus, complicating direct comparisons. Third, the mouse models used vary in the number and identity of triplicated genes, leading to divergent phenotypic expressions. Finally, differences in the nature, source, and dosage of DYRK1A inhibitors may contribute to the observed variability.

In the present study, we investigated the effects of prenatal treatment with Leucettine L41 (L41), a synthetic DYRK1A inhibitor derived from the marine sponge alkaloid Leucettamine B (Debdab *et al*. 2011; Tahtouh *et al*. 2012). L41 has previously demonstrated positive effects in adult DS mice from 3 distinct models (Nguyen *et al*. 2018). We administered L41 to two DS mouse models, Dp1Yey and Ts65Dn, from embryonic day 1 (E1) to delivery. Our objectives were threefold: (1) to evaluate the long-term effects of this standardized treatment on learning and memory at the organismal level; (2) to assess its impact on the density of GAD67 neurons, which influence the excitatory/inhibitory balance; and (3) to examine changes in the expression of brain-derived neurotrophic factor (*Bdnf*), a gene critical for learning and memory (Heldt *et al*. 2007). *BDNF* levels are decreased in the DS brain (Parrini *et al*. 2017) and interventions that increase *Bdnf* expression, such as physical exercise or BDNF-mimetic drugs, have been shown to improve memory in Ts65Dn mice (Parrini et al. 2017). Through a comprehensive characterization of in-utero treated DS mice compared to untreated controls, we aimed to elucidate the differential effects of prenatal L41 treatment across models, likely attributable to their genetic distinctions.

## Materials and Methods

### Mouse model experiments

All experiments were performed following the Directive of the European Parliament: 2010/63/EU, revising/replacing Directive 86/609/EEC and with French Law (Decret n° 2013-118 01 and its supporting annexes entered into legislation 01 February 2013) relative to the protection of animals used in research. YH was the principal investigator of this study (accreditation 67-369) in our animal facility (Agreement C67-218-40). Experimental procedures for the use of animals for research were approved by the Ministry of National Education, Higher Education and Research and with the agreement of the local ethical committee Com’Eth (no. 17) under the accreditation numbers APAFIS #8701-2017012710171063 and #17204-2018102218181793.

The B6EiC3Sn a/A-Ts(17^16^)65Dn/J (RRID:IMSR_JAX:001924; Ts65Dn) mouse line analysed in the study, was obtained from the Jackson Laboratory from the 1924 subline (Shaw *et al*. 2020), and was kept in an F1 B6JC3B genetic background; with the C3B line as a C3H/HeN congenic line for the BALB/c allele at the *Pde6b* locus RRID:IMSR_JAX:013530 (Hoelter et al. 2008). Ts65Dn mice were genotyped according to published protocols (Duchon et al., 2011). The B6.129S7-Dp(16Lipi-Zbtb21)1Yey/J (RRID:IMSR_JAX:013530; Dp1Yey) mice were maintained on a C57BL/6J background and genotyped as described (Li *et al*. 2007). To produce the cohort of animals, we mated B6J females with Dp1Yey males, and Ts65Dn females with F1B6JC3B males. Conception (E0.5, embryonic day 0.5) was determined by examining the vaginal plug. Male and female trisomic and euploid offspring were used for this study.

### Treatment with L41

Leucettine L41 was prepared at 40 mg/ml in dimethyl sulfoxide (DMSO), aliquoted and stored below −20°C (Nguyen *et al*. 2018). The final formulation was prepared just prior to use as a 2 mg/ml solution diluted in PEG300/water (50/45), to reach a final DMSO/PEG300/water 5/50/45 (v/v/v) mix. Pregnant females were randomly assigned to treatment by a daily subcutaneous injection of either L41 (dose: 20 mg/kg/day) or vehicle (the same formulation without L41) from embryonic day 1 to the delivery (embryonic days 20/21). Treated or non-treated pregnant mice were housed individually and were supplied with nesting material. Offspring were kept with their mothers until 3 weeks (21 days) of age, at which time the pups were weaned and separated by sex into cages. From post-natal days 1 (P1) to behaviour assessment, the animals were left without treatment interventions.

### In vivo biodistribution of L41 after injection in pregnant mice

C57BL/6J wildtype (wt, strain 632, Charles River Laboratories, France) females were mated with wt males and pregnant females were injected with unique L41 (20mg per kg) subcutaneous injection at J19 post coitum. The brains and livers of the embryos were collected at different times after L41 subcutaneous injection (10, 20, 30, 60 and 120min) and immediately frozen in liquid azote. Then, each brain and liver lobe were crushed in 400 µL of water and 800 µL of acetonitrile were added to extract the compound. Samples were vortex-stirred for 3 minutes, sonicated for 1 minute and thereafter centrifuged at 16°C, 15,000g during 5 minutes to sediment proteins and suspended tissues. Supernatants were transferred into a microplate and were analysed using a UHPLC coupled to a triple quadrupole. For both kind of samples, standard curves were obtained by analysing known L41 quantities that were dissolved in organ homogenates and analysed using the same procedure.

Analyses were performed on a LCMS 8030 Shimadzu using multiple reaction monitoring mode (MRM). Separations were carried out at 40°C using a 2.6 µm C18 Kinetex column (50 × 2.1 mm) purchased from Phenomenex. The mobile phase flow rate was set at 0.5 mL/min and the following program was applied for the elution: 0 min, 5% B; 0-1.2 min, 5-95% B; 1.2-1.4 min, 95% B; 1.4-1.42 min, 95-5% B and 1.42-2.8 min, 5% B (solvent A: 0.05% formic acid in water; solvent B: acetonitrile). Injection volume was 5 µL. The mass spectrometer was interfaced with the liquid chromatograph using an electrospray ion source. The nitrogen nebulizing gas flow was set at 1.5 L/min and the drying gas flow at 15 mL/min. 4500 V were used for the interface voltage. The temperature of the block heater was maintained at 400 °C and the one of the desolvation line at 250 °C. The collision gas used was nargon at 230 kPa. The MRM transition in positive mode for L41 was m/z 308.0 → 119.2, 160.0 308.0>119.2 and 308.0>160.0.

### DYRK1A kinase activity

C57BL/6J wildtype (wt, strain 632, Charles River Laboratories, France) females were mated with Dp1Yey males and presence of vaginal plug determine the beginning of gestation. At 12, 14, 16 and 18 days of gestation, pregnant females were euthanise and embryo brains were collected and immediately frozen in liquid azote. Then, each brain was lysed with Precellys homogenizer (Bertin Technologies SAS, Montigny-le-Bretonneux FRANCE) in 600 µL of 60 mM β-glycerophosphate, 25 mM Mops (pH 7.2), 15 mM EGTA, 15 mM MgCl2, 2 mM dithiothreitol, 1 mM sodium orthovanadate, 1 mM sodium fluoride, 1 mM phenylphosphate disodium and protease inhibitor buffer. Supernatants were then assayed for protein concentrations with the Pierce BCA Protein Assay Kits (Thermo Fisher Scientific, Waltham, MA USA). To assessed DYRK1A kinase activity, we used a method based on the separation and quantification of specific fluorescent peptides (substrate and phosphorylated product) by LC (Bui *et al*. 2014).

### Behavioural assessment

A series of behavioural experiments were conducted in male mice with an age range starting at 75 up to 120 days for the last test depending on the availability of the test platform. We selected male mice only to reduce the number of animal use in research as we know that L41 treatment can rescue cognition in both sexes in adults (Nguyen *et al*. 2018). After weaning, animals were sorted by litters into 39 × 20 × 16 cm cages (Green Line, Techniplast, Italy) where they had free access to purified water and food (D04 chow diet, Safe, Augy, *France*). The temperature was maintained at 23±1 °C, and the light cycle was controlled as 12 h light and 12 h dark (lights on at 7 am). On testing days, animals were transferred to the antechambers of the experimental room 30 min before the start of the experiment. All experiments were performed between 8:00 AM and 4:00 PM. The tests were administered in the following order: week 1, nesting, and week 2 Y-maze, open field and novel object recognition.

### Nesting test

The nesting test was performed by placing the mice individually in clean new housing cages 2 h before the dark phase, and the results were assessed the following day (Deacon 2012). Regular bedding covered the floor to a depth of 0.5 cm. Each cage had a ‘nestlet’, a 5 cm square of pressed cotton batting. The nests were assessed on a five-point scale: 1, the nestlet was largely untouched (>90% intact); 2, the nestlet was partially torn up (50-90% remaining intact); 3, the nestlet was mostly shredded but often there was no identifiable nest site: <50% of the nestlet remained intact but <90% was within a quarter of the cage floor area, i.e. the cotton was not gathered into a nest but spread around the cage; 4, an identifiable but flat nest: >90% of the nestlet was torn up; the material was gathered into a nest within a quarter of the cage floor area, but the nest was flat with walls higher than mouse body height (curled up on its side) on less than 50% of its circumference; 5, a (near) perfect nest: >90% of the nestlet was torn up; the nest was a crater, with walls higher than mouse body height for more than 50% of its circumference (Deacon 2012). For statistical analysis, the score was transformed in binomial data like 1 for score 4 or 5 (present of nest) and 0 for score 1, 2 or 3 (absence of nest).

### Y-maze

Short-term working memory was assessed by recording spontaneous alternation in the Y-maze test (Hughes, 2004). The Y-maze test is based on the innate preference of animals to explore an arm that has not been explored previously, a behaviour that, if occurring with a frequency greater than 50%, is called spontaneous alternation behaviour (SAB). The maze was made of three enclosed plastic arms, each 40×9×16 cm, set at an angle of 120° to each other in the shape of a ‘Y’. The wall of each arm had a different pattern to encourage SAB. Animals were placed at the end of one arm (this initial arm was alternated within the group of mice to prevent arm placement bias), facing away from the centre, and allowed to freely explore the apparatus for 8 min under moderate lighting conditions (70 lux in the centre most region). The time sequences of entries in the three arms were recorded (considering that the mouse entered an arm when all four paws were inside the arm). Alternation was determined from successive entries into the three arms on overlapping triplet sets in which three different arms are entered. The number of alternations was then divided by the number of alternation opportunities, namely, total arm entries minus one. In addition, total entries were scored as an index of locomotor activity.

### Open field

The open-field test (OF) measures locomotor activity, exploratory drive and some aspects of anxiety in mice. The output of the various interacting drives is locomotion, which is the direct measure obtained. The OF was performed in a white circular arena (55-cm diameter) placed in a dimly lit testing room (40 lux). The mice were introduced for two sessions of 15 minutes with 24 hours interval. During those sessions, mice were monitored using a video tracking system (Ethovision, Wageningen, The Netherlands), and the total distance travelled in the different zones of the arena (centre, intermediate and peripheral) was recorded.

### Novel object recognition test

The Novel object recognition (NOR) task is based on the innate tendency of rodents to explore novel objects over familiar ones (Bevins and Besheer, 2006). This test was done 24 h after the last OF session performed in the same arena. On day 1, mice were free to explore two identical objects for 10 min. After this acquisition phase, mice returned to their home cage for a 24 h retention interval. Their memory was evaluated on day 2 in the test session, using one familiar object (of those already experienced during the acquisition phase) and one novel object, which were placed in the arena with the mice free to explore the two objects for 10 minutes. The discrimination index (DI=(time of exploration of the new object – time of exploration of the familiar object) / time of total exploration) therefore measured their memory performance. The closer the index was to zero, the worse the memory. This was tested statistically by a one-sample t-test versus 0. Between trials and subjects, the different objects were cleaned with 70% ethanol to reduce olfactory cues. To avoid any preference for one of the two objects or their relative location, object type and location were randomly assigned and balanced for each individual. Object exploration was manually scored and defined as the orientation of the nose to the object, allowing a distance <1 cm. Activity in the test was evaluated by the distance travelled by the mice in the arena.

### Bdnf gene expression by quantitative real time polymerase chain reaction (RT-qPCR) analysis

Hippocampi were isolated from DS trisomic models and their littermate controls and flash-frozen in liquid nitrogen. Total RNA was prepared using the RNA extraction kit (Qiagen, Venlo, The Netherlands) according to the manufacturer’s instructions. cDNA synthesis was performed using the SuperScript® VILO™ cDNA Synthesis Kit (Invitrogen, Carlsbad, CA). PCR were performed with TaqMan® Universal Master Mix II and pre-optimized TaqMan® Gene Expression assays (Applied Biosystems, Waltham, Massachusetts, USA), consisting of a pair of unlabeled PCR primers and a TaqMan® probe with an Applied Biosystems™ FAM™ dye label on the 5’ end and minor groove binder (MGB) and nonfluorescent quencher (NFQ) on the 3’ end. mRNA expression profiles were analysed by real-time quantitative PCR using TaqMan TM universal master mix II with UNG in a realplex II master cycler Eppendorf (Hambourg, Germany). The complete reactions were subjected to the following program of thermal cycling: 1 cycle of 2 minutes at 50°C, 1 cycle of 10 minutes at 95°C, 40 cycles of 15 seconds at 95°C and 1 minute at 60°C. Efficiencies of the TaqMan assays were checked using cDNA dilution series from extracts of hippocampal sample. Normalization was performed by carrying out in parallel the amplification of 4 housekeeping genes (*Gnas, Pgk1, Actb* and *Atp5b*) and using the GeNorm procedure to correct the variations of the amount of source RNA in the starting material (Vandesompele *et al*. 2002). All the samples were tested in triplicate.

### Histological procedures

Mice received an intra-peritoneal overdose of ketamine (150mg/kg) / xylasine (10 mg/kg) and were perfused transcardially with 20ml of phosphate-buffered followed by 30ml of 4% phosphate-buffered paraformaldehyde (PFA). Brains were removed, post-fixed overnight (4% PFA, 4°C), and transferred to PBS for 2 days with 2 medium changes. A cryostat cut the complete brains extending over the entire hippocampus into 64 free-floating coronal sections of 50 μm. These sections were extended from +4.00 to +2.52 mm from Bregma, i.e. 3.2mm. All tissue sections were collected and stored in PBS at 4°C before being processed. Every 6th section was sub-sampled, starting randomly between section 1 to 6, for final immunohistochemistry and stereological quantification.

For Immunohistochemistry, endogenous peroxidases were first quenched with 0.3% H2O2 for 30 min. After washes, slices were incubated with blocking solution for 1 hour at RT. After washes, incubation with the primary antibody overnight at 4°C was done followed by incubation of the secondary antibody (biotinylated) in for 1 h at RT. We used primary mouse anti-GAD67 MAB5406, rabbit anti-NEUN ZRB377 (Sigma Aldrich Chimie S.a.r.l, Saint-Quentin-Fallavier, France). Secondary antibodies were rabbit Anti-Goat IgG antibody (H+L) biotinylated and goat Anti-mouse IgG antibody (H+L) Biotinylated (Vector Laboratories, Burlingame, CA). Finally, Kit ABC Vectastain and impact DAB peroxidase substrate (Vector Laboratories, Burlingame, CA) was used for peroxidase revelation.

Finally, sections of interest were digitized with a slide scanner (Hamamatsu Nanozoomer 2.0HT) equipped with a Z-stack feature that allows to focus on different depths in the sample. Then, analysis of hippocampus stack images was performed using the NIH ImageJ software version 4.7. The counting of cells was done in zones covering stratum radiatum, stratum lacunosum-moleculare, and stratum moleculare because of the presence of GABAergic interneurons in this era (Klausberger and Somogyi 2008). The optimal number, size and spacing of optical dissectors were determined to obtain a coefficient of error (CE) under 0.10 (West 1990). For unbiased stereological assessment, the counting parameters were defined as follows: For NEUN, side dissector size = 91.0996 microns, height of optical dissector: 20µm, area of counting frame = 8299.132 µm2 and area of grid spacing = 27387.1356 µm2. For GAD67, side dissector size = 136.6 microns, height of optical dissector: 20µm, area of counting frame = 18671.34 µm2 and area of grid spacing = 40330.10 µm2. The estimated cell number was calculated with the formula: cells counted x Xth series x (area of grid spacing / area of counting frame) x (mean measured thickness of section / height of optical dissector).

### Statistical analysis

The accuracy of a statistical analysis is largely dependent on the size of the sample, a sample size too small may not be representative of the population as a whole, leading to inaccurate conclusions (Faber and Fonseca 2014). In the present study, the number of individuals sexes groups was too low to have robust conclusion, so we don’t assessed sexes effects, female and male data were pooled. Experiments were carried out using a double-blind approach, namely persons in charge of the recording of behaviour sessions and of the quantitative analyses did not know about the treatments of the experimental groups. In addition, wild-type and Dp1Yey, or Ts65Dn, females with plug were assigned, at random, to vehicle or L41-injected groups. For nesting (Figure 2), score was transformed in binary data. Cohen’s h (measure of distance between two proportions) and the corresponding proportion plot were produced (Ho et al. 2019). 5000 bootstrap samples were taken; the confidence interval is bias-corrected and accelerated. Any p-value reported is the probability of observing the effect size (or greater), assuming the null hypothesis of zero difference is true. For each p-value, 5000 reshuffles of the control and test labels were performed.

For dataset coming from behaviour tests, we performed the Shapiro–Wilk test and quantile-quantile plots to analyse whether the data were normally distributed and the Brown–Forsythe test to ascertain the homogeneity of variances. If the P-value was greater than the significance level (0.05), we assumed normality and equal variance. In this case, the statistical significance of differences between groups was inferred by an ANOVA genotype treatment. The post hoc tests (Tukey test) were conducted only if F in repeated measures ANOVA achieved a 0.05 level. In the case of datasets in which the assumptions of normality or homogeneity of variances were not fulfilled, we used an ANOVA on the rank non-parametric test, with Dunn’s Method for all pairwise multiple comparison procedures. For Ymaze and NOR analysis, we performed in addition a one-sample two-tailed t-test for the percentage of spontaneous alternation versus 50% (hazard) or the DI versus no discrimination (0%).

In the cases where the data have less than n<=8 in one parameter, statistical differences were assessed by a permutation test. This is the case for qPCR relative expression, immunohistological, L41 biodistribution and kinase activity assay (Figure 6**Error! Reference source not found**., Figure, Figure 7 and **Error! Reference source not found**.). To analyse these data, we used DABEST (Data Analysis with Bootstrap-coupled ESTimation), open-source libraries for Python, to calculate the 95% confidence interval of the mean difference for treatment or genotype by performing bootstrap resampling (5000) with bias-corrected and accelerated bootstrap correction. This enables visualization of the confidence interval as a graded sampling distribution. If 0 is out of the 95% confidence interval of the effect size (Δ), the null hypothesis is rejected with pValue < 0.05. There is no need to assume that observations are normally distributed. Thanks to the Central Limit Theorem, the resampling distribution of the effect size will approach normality. Results are presented as Gardner-Altman estimation plot. All data points were presented as a swarmplot, the effect size was presented as a bootstrap 95% confidence interval (Ho et al. 2019). Completes results are presented in **Error! Reference source not found**..

## RESULTS

### Biodistribution of Leucettine L41 in the Embryonic Brain after Maternal Administration

For prenatal treatment to be effective, the administered compound must successfully traverse both the placental barrier and the developing blood-brain barrier to reach the embryonic brain. To evaluate this critical requirement, wild-type females were mated with wild-type males, and pregnant dams at embryonic day 19 (E19) received a single subcutaneous injection of L41.

The biodistribution of L41 systematically assessed in vivo at multiple time points following administration. Quantitative analysis demonstrated the presence of detectable L41 levels in embryonic tissues, with a peak concentration observed at 30 minutes post-injection (Figure 1A). These findings confirm that L41 effectively crosses both the placental and embryonic blood-brain barriers, thereby validating its potential for prenatal therapeutic intervention.

**Figure 1:**
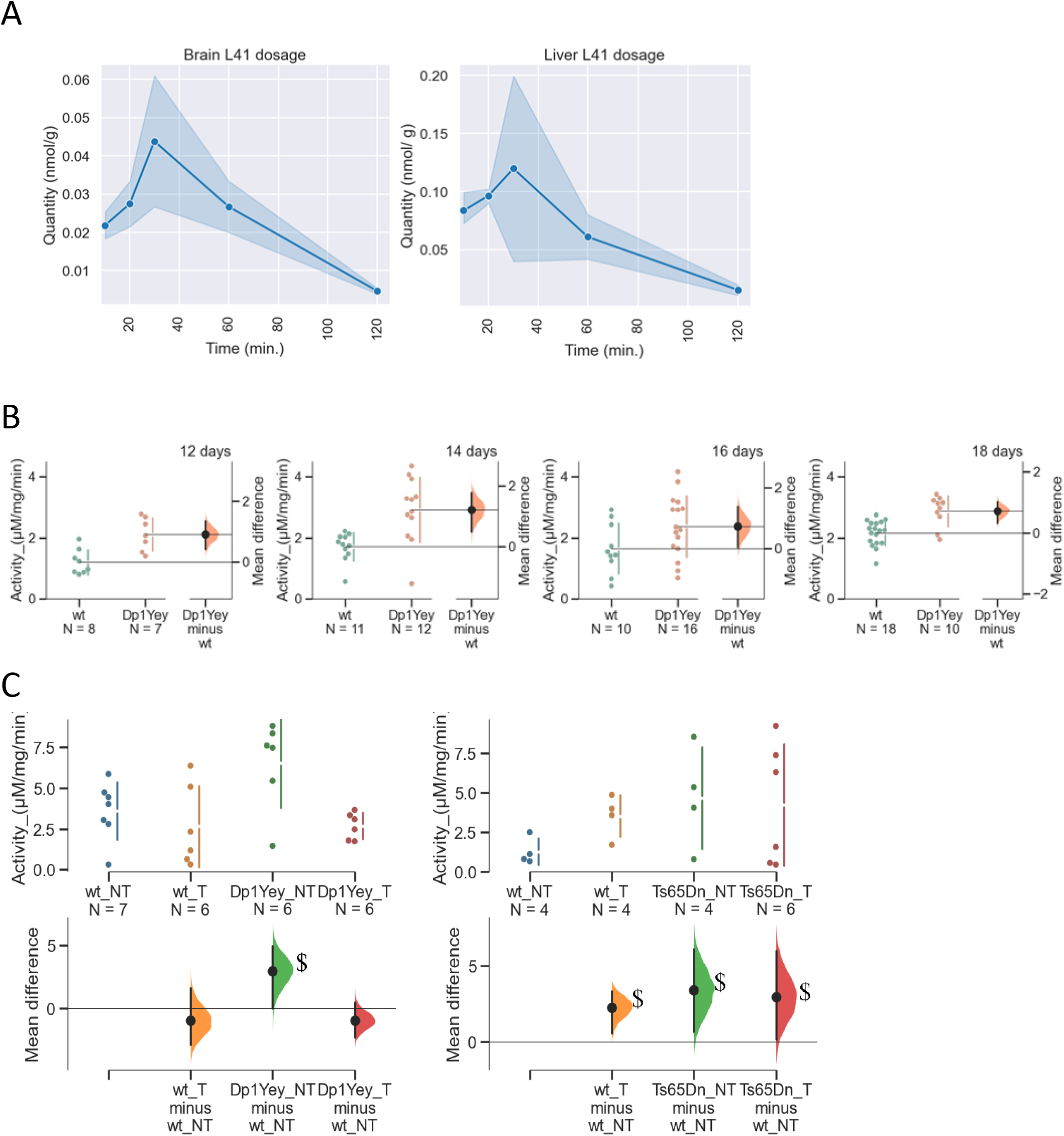
A single prenatal injection of L41 reduces DYRK1A kinase activity in the brains of E19 trisomic embryos. (A) Time-course quantification of L41 concentration in embryonic brain and liver following a single subcutaneous injection at embryonic day 19 (E19). Data are shown for 2 to 4 embryos per time point. (B) DYRK1A kinase activity measured at embryonic days E12, E14, E16, and E18. Results are presented as Gardner-Altman estimation plots. Left panels: Raw activity values for each group. Right panels: Bootstrap sampling distributions of the mean differences between treated and untreated groups (wild-type and trisomic). Black dots represent the observed mean differences. Horizontal black lines indicate the 95% confidence intervals based on 5,000 bootstrap resamples. The symbol $ indicates that zero is outside the 95% CI, denoting a statistically significant difference. (C) DYRK1A activity (expressed in µg/mg/min) in E19 embryonic brains after a single L41 injection. In untreated trisomic embryos, DYRK1A activity was significantly elevated compared to wild-type. L41 treatment reduced this activity to wild-type levels in the Dp1Yey model. No significant treatment effect was observed in Ts65Dn embryos, likely due to high inter-individual variability. Data are also shown as Gardner-Altman plots, with raw values in the upper panels and bootstrap distributions of mean differences in the lower panels. As above, $ indicates that zero lies outside the 95% CI.

**Figure 2:**
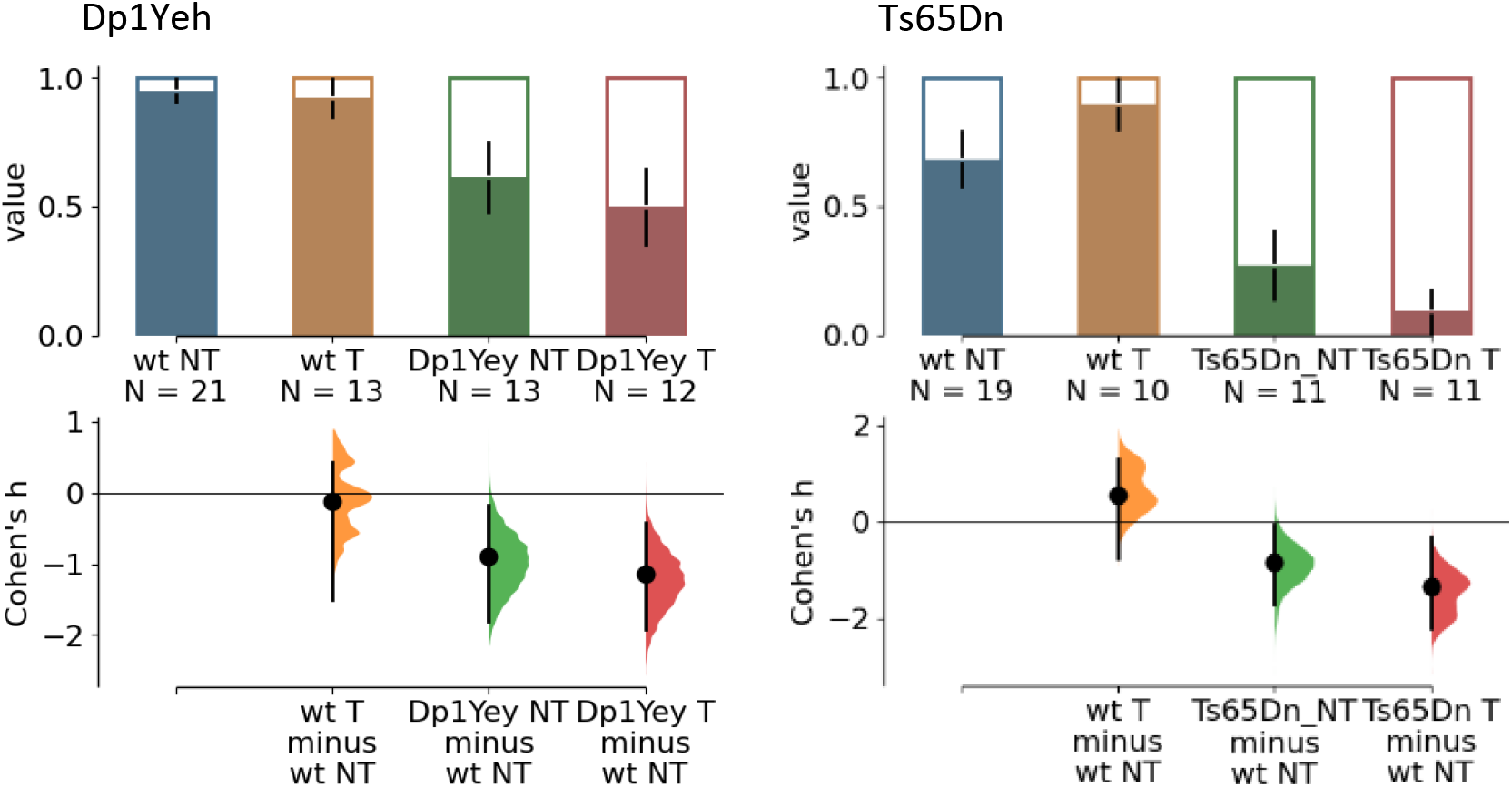
Nesting activity is not improved after L41-prenatal treatment in the adult progeny. DS mouse models mice that made a nest (value 1) and the white part in the bar represents the proportion of mice that did not do a nest (value 0) in the dataset. Cohen’s h and the corresponding proportion plot for binary data were produced. 5000 bootstrap samples were taken; the confidence interval is bias-corrected and accelerated.

### Elevated DYRK1A Kinase Activity in the Developing Brain of Dp1Yey Mouse Embryos

We hypothesized that increased DYRK1A kinase activity during gestation contributes to the neurodevelopmental alterations observed in trisomic embryos. To test this, we quantified DYRK1A activity in brain protein extracts from Dp1Yey and wild-type embryos at four key developmental stages: embryonic days E12, E14, E16, and E18. Our results confirmed a consistent and significant elevation of DYRK1A activity in Dp1Yey embryos relative to wild-type controls across all examined time points (E12: 95% CI [0.42, 1.35]; E14: [0.48, 1.77]; E16: [0.04, 1.38]; E18: [0.31, 1.04]; Figure 1B). These findings support the role of DYRK1A overexpression in the pathogenesis of neurodevelopmental phenotypes associated with Down syndrome.

### Differential Efficacy of Prenatal L41 Treatment on DYRK1A Activity in Dp1Yey and Ts65Dn Embryos

To evaluate the capacity of L41 to mitigate DYRK1A overactivity, pregnant dams carrying either Dp1Yey or Ts65Dn embryos received a single subcutaneous injection of L41 at a dose of 20 mg/kg. Embryos were collected at embryonic day 18 (E18), one hour post-injection, and brain tissues were analyzed for DYRK1A kinase activity. In untreated embryos, DYRK1A activity was significantly elevated in both trisomic models relative to wild-type controls (Dp1Yey: 95% CI [0.01, 4.90]; Ts65Dn: [0.66, 6.14]; Figure 1C).

Following L41 administration, DYRK1A activity in Dp1Yey embryos was effectively reduced to levels indistinguishable from those of wild-type controls (Dp1Yey treated vs. wild-type untreated: 95% CI [–2.27, 0.49]), demonstrating successful inhibition of kinase overactivity. However, L41 treatment failed to produce a significant reduction in DYRK1A activity in Ts65Dn embryos. This lack of effect may be attributed to substantial inter-individual variability, as Ts65Dn embryos appeared to segregate into two distinct subgroups: one exhibiting a robust response to treatment and the other showing minimal to no response (Figure 1C). Such heterogeneity could arise from differences in pharmacokinetics, blood brain barrier penetration, genetic background, or compensatory regulatory mechanisms within this model.

### Prenatal L41 Treatment Selectively Improves Survival Rates in Ts65Dn but Not Dp1Yey Down Syndrome Mouse Models

To generate the required cohorts, B6N females were mated with Dp1Yey males, and Ts65Dn females were mated with F1B6C3B males. Fertilization was confirmed by the detection of a vaginal plug, marking the initiation of treatment with either L41 or a vehicle control. For the Dp1Yey line, 21 untreated (NT) females and 12 treated (T) females were used, resulting in pregnancy rates of 86% (18/21) and 83% (10/12), respectively. L41 treatment did not affect pregnancy rates in wild-type (WT) females mated with Dp1Yey males.

Dp1Yey matings produced a total of 165 pups: 52 live pups in the NT group (38 WT, 14 Dp1Yey) and 40 live pups in the T group (28 WT, 12 Dp1Yey; Table 1). Chi-square analysis revealed that pup mortality was significantly associated with genotype in both the NT group (χ^2^(1, N = 103) = 10.03, p = 0.002) and the T group (χ^2^(1, N = 61) = 4.26, p = 0.039), but not with treatment. For behavioral testing, cohorts were selected as follows: 21 untreated WT (9 females, 12 males), 13 treated WT (6 females, 7 males), 13 untreated Dp1Yey (6 females, 7 males), and 12 treated Dp1Yey (3 females, 9 males).

**Table 1:** Viability of the progeny from untreated (NT) or treated (T) pregnant females from Dp1Yey and Ts65Dn DS mouse lines. Chi square test, *p < 0.05, **p < 0.01, ***p < 0.001.

| Treatment | Crosses | B6.Dp(16)1Yey male X WT B6 Female |  |  | B6C3B WT male X B6C3B Ts65Dn female |  |  |
| --- | --- | --- | --- | --- | --- | --- | --- |
|  | Viable at birth | WT | Dp1Yey | Total | WT | Ts65Dn | Total |
| NT | Yes | 38 | 13 | 51 | 32 | 12 | 44 |
| NT | No | 23 | 29 | 52 | 35 | 33 | 68 |
| NT | Total | 61 | 42 | 103 | 67 | 45 | 112 |
|  | % of death at birth | 38 | 67 | ** | 52 | 73 | * |
| T | Yes | 28 | 12 | 40 | 10 | 11 | 21 |
| T | No | 9 | 12 | 21 | 8 | 6 | 14 |
| T | Total | 37 | 24 | 61 | 18 | 17 | 35 |
|  | % of death at birth | 24 | 50 | * | 44 | 35 | n.s. |

For the Ts65Dn model, 25 NT females and 11 T females were used, resulting in pregnancy rates of 68% (17/25) and 55% (6/11), respectively. The pregnancy rates were not statistically different between the Dp1Yey and Ts65Dn lines following L41 treatment. Ts65Dn matings yielded 147 pups: 44 live pups in the NT group (32 WT, 12 Ts65Dn) and 21 live pups in the T group (10 WT, 11 Ts65Dn; Table 2). Genotype significantly influenced pup mortality in the NT group (χ^2^(1, N = 104) = 4.72, p = 0.03), but not in the T group (χ^2^(1, N = 35) = 0.30, p = 0.58), suggesting a potential survival benefit of L41 treatment in Ts65Dn pups. Behavioral testing cohorts included 19 untreated WT (10 females, 9 males), 10 treated WT (4 females, 6 males), 11 untreated Ts65Dn (6 females, 5 males), and 11 treated Ts65Dn (4 females, 7 males).

Behavioral assessments were conducted on these animals, which had been treated prenatally with L41 at a dose of 20 mg/kg. Testing commenced at 90 days post-partum to evaluate the long-term effects of in-utero L41 administration on adult mice.

### Prenatal L41 Treatment Fails to Rescue Nesting Deficits in Dp1Yey and Ts65Dn Down Syndrome Mouse Models

Nesting behavior was evaluated by scoring the quality of nests constructed overnight from a standardized cotton sample (Deacon 2012). In the Dp1Yey model (Figure 2), untreated Dp1Yey built nests of significantly lower quality compared to WT controls (Cohen’s h = –0.898, 95% CI [–1.80, – 0.179], p = 0.046, two-sided permutation t-test). However, prenatal L41 treatment did not improve nesting performance: the difference between WT NT and Dp1Yey T groups remained significant (Cohen’s h = –1.13, 95% CI [–1.91, –0.42], p = 0.0258), indicating that L41 had no detectable effect on this behavior. A similar pattern was observed in the Ts65Dn model (Figure 2). Untreated Ts65Dn mice exhibited impaired nesting compared to WT controls (Cohen’s h = –0.849, 95% CI [–1.71, –0.0675], p = 0.0266). L41 treatment likewise failed to rescue this deficit, as the difference between WT NT and Ts65Dn T groups remained significant (Cohen’s h = –1.34, 95% CI [–2.19, –0.313], p = 0.0004). These findings indicate that while both Dp1Yey and Ts65Dn mice display impaired nesting behavior, prenatal L41 treatment does not ameliorate this phenotype in either model.

### Prenatal L41 Treatment Rescues Working Memory Deficits in Adult Dp1Yey Mice but Fails to Improve Performance in Ts65Dn Mice

Working memory was assessed using the Y-maze spontaneous alternation task, which evaluates short-term spatial memory by measuring the tendency of rodents to explore new arms rather than revisit previously visited ones. During a 6-minute session, locomotor activity was quantified by the number of arm entries, while working memory performance was assessed by the percentage of spontaneous alternations.

In the Dp1Yey model, untreated mice exhibited significantly increased locomotor activity compared to wild-type controls (F(1,55) = 19.90, p < 0.001; Figure 3), with no significant effect of L41 treatment on activity levels (F(1,55) = 3.39, p = 0.071). However, Dp1Yey mice displayed a marked deficit in spontaneous alternation (F(1,55) = 9.68, p = 0.003). Importantly, prenatal L41 treatment significantly improved alternation performance in Dp1Yey mice, indicating a restoration of working memory (genotype × treatment interaction: F(1,55) = 3.39, p = 0.031).

**Figure 3:**
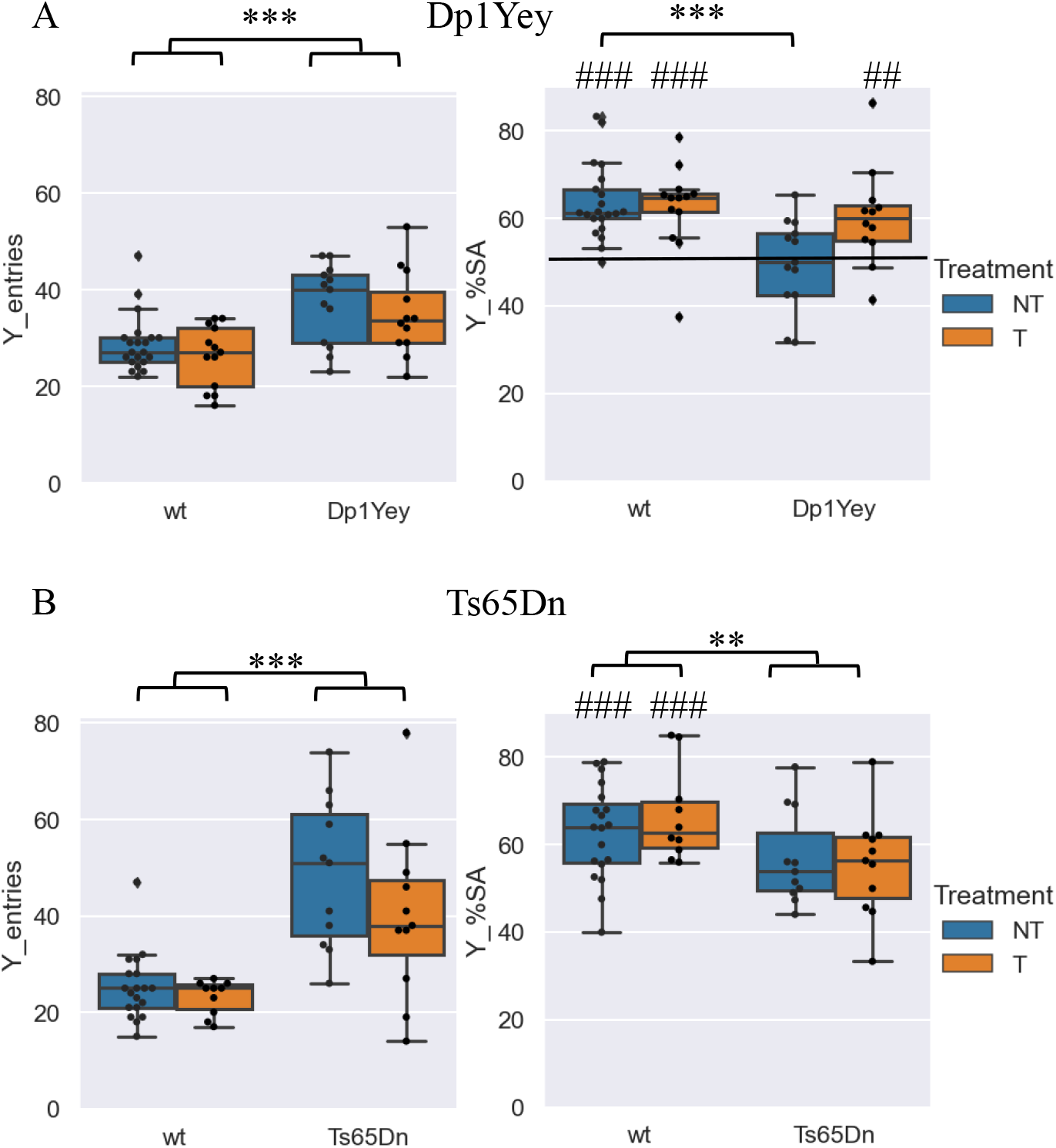
Locomotor activity and spontaneous alternation performance in the Y-maze following prenatal L41 treatment in Dp1Yey and Ts65Dn mice. Box plots represent the number of arm entries (left panels) and the percentage of spontaneous alternation (right panels) for each genotype and treatment group. Boxes indicate the interquartile range (IQR), with the median shown as a horizontal line; whiskers represent the 90% confidence interval. (A) Dp1Yey model: Dp1Yey mice exhibited significantly increased locomotor activity compared to wild-type (wt) controls, with no effect of L41 treatment. A significant reduction in spontaneous alternation was observed in untreated Dp1Yey mice compared to wt. However, L41 treatment restored alternation performance to wild-type levels. One-sample t-tests confirmed that alternation rates in wt NT, wt T, and Dp1Yey T groups were significantly above chance level (50%), whereas Dp1Yey NT mice did not differ from chance. These results indicate that prenatal L41 treatment rescues working memory deficits in Dp1Yey mice. (wt NT n = 21; wt T n = 13; Dp1Yey NT n = 13; Dp1Yey T n = 12). (B) Ts65Dn model: Ts65Dn mice also showed hyperactivity, with no significant effect of treatment. A significant genotype effect was observed on spontaneous alternation, with Ts65Dn mice performing below wt levels. However, L41 treatment did not improve alternation performance. One-sample t-tests confirmed that alternation rates in Ts65Dn groups remained at or below chance level. (wt NT n = 19; wt T n = 10; Ts65Dn NT n = 11; Ts65Dn T n = 11). Statistical significance: ANOVA p < 0.01 (**), p < 0.001 (***); one-sample t-test vs. 50%: # p < 0.05, ## p < 0.01, ### p < 0.001.

In contrast, Ts65Dn mice demonstrated pronounced hyperactivity (Kruskal–Wallis test: H(3) = 23.07, p < 0.001), with no significant effect of L41 treatment on locomotor activity. These mice also exhibited a significant reduction in spontaneous alternation compared to wild-type controls (F(1,47) = 7.48, p = 0.009). However, unlike in the Dp1Yey model, L41 treatment did not improve alternation performance in Ts65Dn mice (F(1,47) = 0.13, p = 0.725; Figure 3).

Thus, prenatal L41 treatment selectively rescued working memory deficits in the Dp1Yey model but not in Ts65Dn mice, underscoring potential model-specific differences in responsiveness to DYRK1A inhibition.

### Prenatal L41 Treatment Does Not Affect General Locomotor Activity During the Habituation Phase of the Novel Object Recognition Test

General locomotor activity was assessed by measuring the distance travelled in an open field during the habituation phase of the Novel Object Recognition (NOR) test. The objective was to familiarize the animals with the testing environment, a circular arena placed in a low-light room. Mice were exposed to the empty arena for two 15-minute sessions (S1 and S2), spaced 24 hours apart. A video tracking system recorded the total distance travelled during each session.

In the Dp1Yey model, mice exhibited significantly increased locomotor activity during Session 1 compared to wild-type controls (*H*(3) = 29.26, *p* < 0.001; Dunn’s post hoc test: WT NT vs. Dp1Yey NT, *p* < 0.001; Figure 4). L41 treatment had no effect on activity levels (Dp1Yey NT vs. Dp1Yey T, *p* = 1.0, n.s.). By Session 2, Dp1Yey mice had habituated to the environment, and their activity levels decreased to match those of wild-type mice (*H*(3) = 0.78, *p* = 0.854, n.s.).

**Figure 4:**
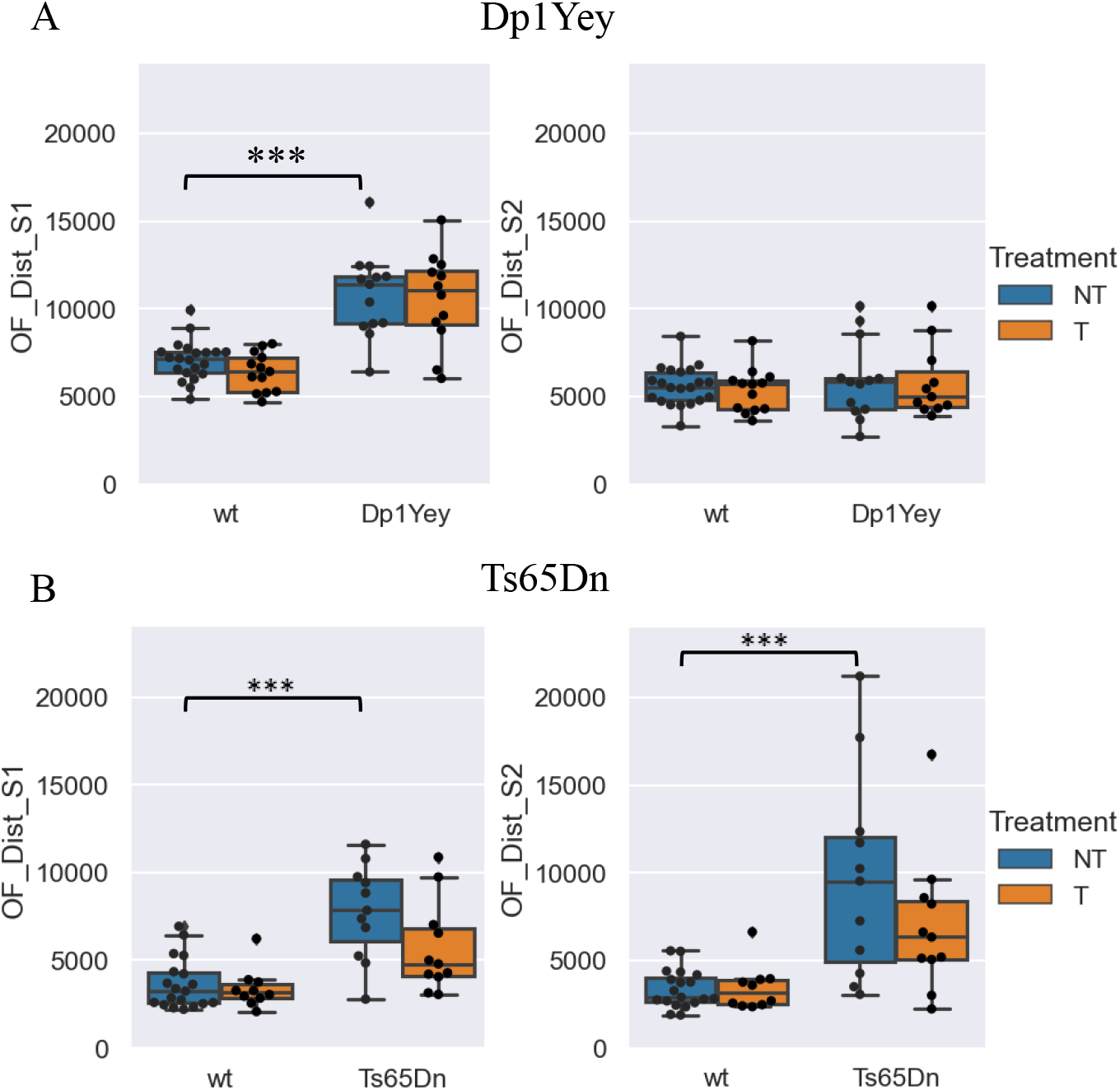
Locomotor activity measured as distance travelled during two consecutive open field sessions (S1 and S2) in Dp1Yey (A) and Ts65Dn (B) mouse models of Down syndrome, with or without prenatal L41 treatment. Box plots represent the total distance travelled during each session, with medians and interquartile ranges; whiskers indicate the 90% confidence interval (grey shading). (A) In session 1, both untreated and treated Dp1Yey mice exhibited significantly increased activity compared to wild-type controls. All groups showed reduced activity in session 2, indicating habituation to the testing environment. (wt NT n = 21; wt T n = 13; Dp1Yey NT n = 13; Dp1Yey T n = 12). (B) In contrast, Ts65Dn mice displayed persistent hyperactivity across both sessions, with no significant reduction in activity between S1 and S2, suggesting impaired habituation. Prenatal L41 treatment had no measurable effect on locomotor activity in either model. (wt NT n = 19; wt T n = 10; Ts65Dn NT n = 11; Ts65Dn T n = 11). Statistical significance: p < 0.05 (*), p < 0.01 (**), p < 0.001 (***).

In the Ts65Dn model, a similar genotype effect was observed in Session 1 (*H*(3) = 20.36, *p* < 0.001; WT NT vs. Ts65Dn NT, *p* < 0.001), with no significant effect of L41 treatment (Ts65Dn NT vs. Ts65Dn T, *p* = 1.0; Figure 4). Unlike Dp1Yey mice, Ts65Dn mice failed to habituate in Session 2, maintaining elevated activity levels (*H*(3) = 20.45, *p* < 0.001; WT NT vs. Ts65Dn NT, *p* < 0.001), with no measurable effect of L41 treatment (Ts65Dn NT vs. Ts65Dn T, *p* = 1.0; Figure 4).

These findings demonstrate that while Dp1Yey mice adapt to the testing environment over time, Ts65Dn mice exhibit persistent hyperactivity. Importantly, prenatal L41 treatment did not modulate general locomotor activity in either model.

### Prenatal L41 Treatment Selectively Restores Episodic Memory Deficits in Trisomic Mice

Long-term episodic memory was evaluated using the Novel Object Recognition (NOR) test. During the initial session (S1, “presentation phase”), mice were placed in an arena containing two identical objects, and the time spent exploring each object was recorded to assess exploratory behavior and object interaction. While Ts65Dn mice exhibited pronounced hyperactivity across both sessions, this did not interfere with object exploration, as no significant differences in interaction time were observed between genotypes or treatment groups.

In the untreated groups, both Dp1Yey and Ts65Dn mice displayed discrimination indices (DI) near zero (Figure 5), indicating impaired recognition memory compared to wild-type controls (one-sample t-test vs. 0: WT NT, t(19) = 3.436, p = 0.0027; WT T, t(11) = 5.607, p < 0.001; Dp1Yey NT, t(13) = 0.534, p = 0.603). Conversely, trisomic mice that received prenatal L41 treatment demonstrated significantly improved DI scores (Dp1Yey T, t(10) = 3.529, p = 0.003; Ts65Dn T, t(10) = 3.001, p = 0.0133), indicating a restoration of recognition memory.

**Figure 5:**
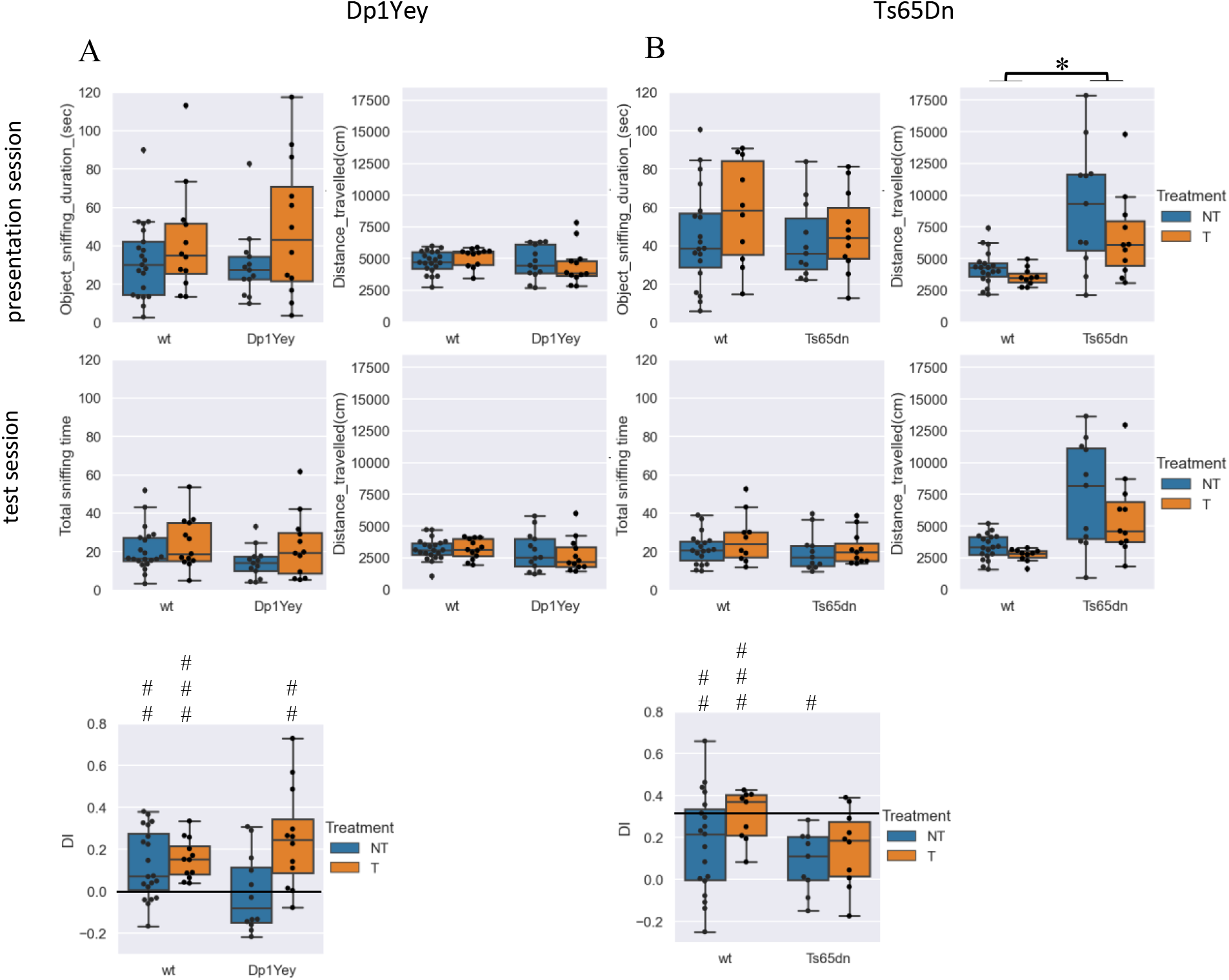
Prenatal L41 treatment restores episodic memory in Dp1Yey (A) and Ts65Dn (B) mouse models of Down syndrome. The panels show (from top to bottom): total sniffing time (in seconds), distance travelled (in cm) during the presentation and test sessions of the Novel Object Recognition (NOR) task, and the discrimination index (DI) reflecting memory performance. Data are presented as box plots showing the median and interquartile range; whiskers represent the 90% confidence interval (grey shading). The Sample sizes were for Dp1Yey panel (A): wt NT n = 20, wt T n = 12, Dp1Yey NT n = 13, Dp1Yey T n = 11; and for Ts65Dn panel (B): wt NT n = 19, wt T n = 10, Ts65Dn NT n = 11, Ts65Dn T n = 11. In both models, prenatal L41 treatment significantly improved the DI in trisomic mice, restoring performance to levels comparable to wild-type controls. No significant differences in object exploration or locomotor activity were observed between groups, indicating that memory improvement was not due to altered exploration behaviour. Statistical significance: Asterisks indicate group comparisons (p < 0.05 (*), p < 0.01 (**), p < 0.001 (***)); Hash symbols indicate one-sample t-tests vs. chance level (DI = 0): # p < 0.05, ## p < 0.01, ### p < 0.001.

**Figure 6:**
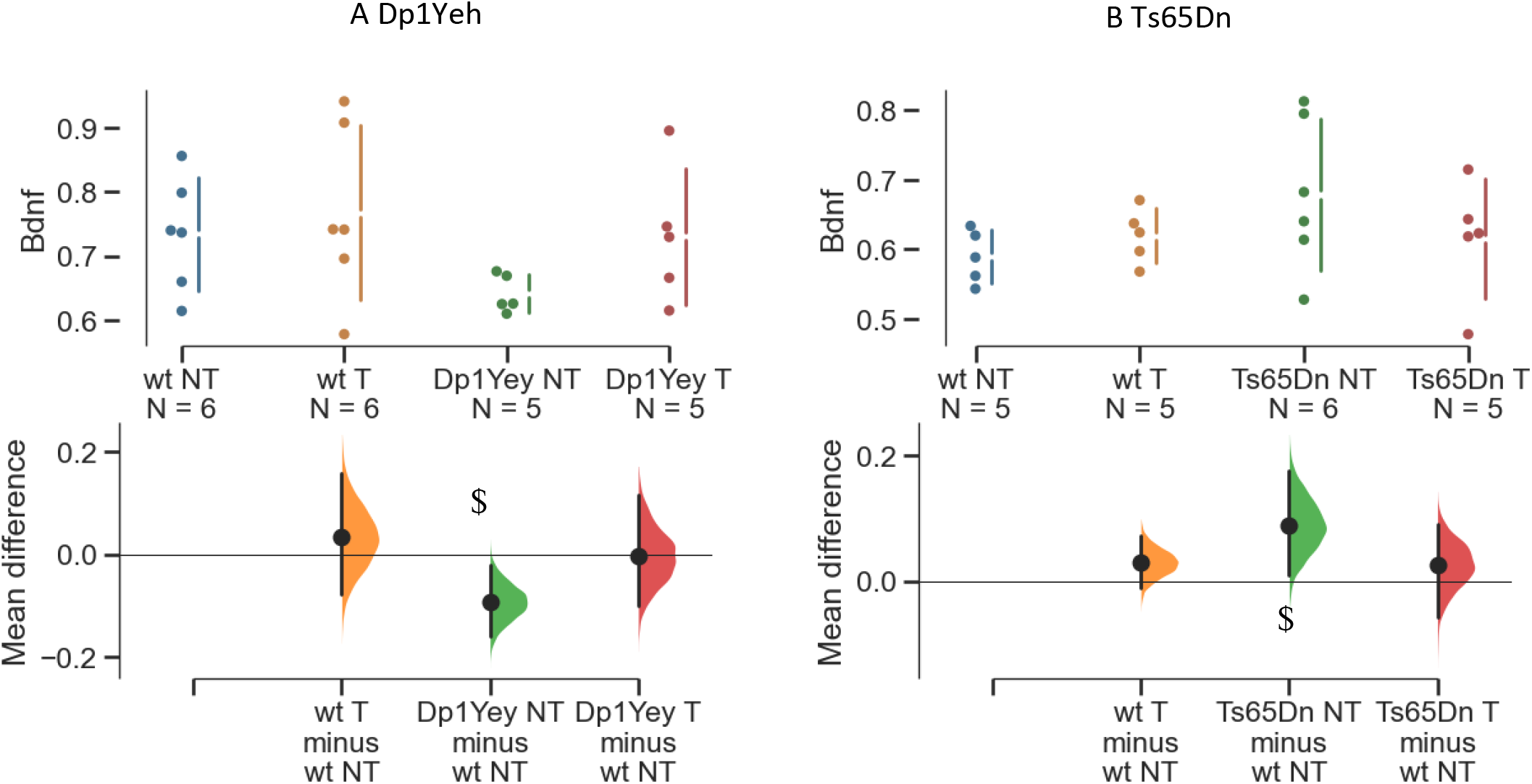
Relative expression levels of Bdnf in the hippocampus of Dp1Yey (A) and Ts65Dn (B) mice, with or without prenatal L41 treatment. The effect of L41 treatment on *Bdnf* expression observed in both wild-type and trisomic mice, highlighting the normalization of expression levels in treated trisomic animals. Data are presented using Gardner-Altman estimation plots. Upper panels: Raw individual data points for each group are shown, allowing visualization of distribution and variability. Lower panels: The bootstrap sampling distribution of the mean difference between treated and untreated groups is displayed. Black dots represent the observed mean differences. Horizontal black lines indicate the 95% confidence intervals, calculated from 5,000 bootstrap resamples.

Notably, L41 treatment did not affect the performance of wild-type mice, suggesting that the in-utero intervention specifically enhances memory function in trisomic models without altering baseline cognition (Figure 5). These findings highlight the potential of prenatal DYRK1A inhibition as a targeted therapeutic strategy for improving episodic memory deficits in Down syndrome.

### Prenatal L41 Treatment Restores Hippocampal Bdnf Expression to Wild-Type Levels in Trisomic Mouse Models

Brain-derived neurotrophic factor (BDNF) plays a critical role in memory formation and synaptic plasticity (Heldt *et al*. 2007)). To investigate its involvement in Down syndrome (DS) pathology, we examined *Bdnf* gene expression in the hippocampus of two trisomic mouse models: Dp1Yey and Ts65Dn.

In Dp1Yey mice, *Bdnf* expression was significantly downregulated compared to WT (WT NT vs. Dp1Yey NT, 95% CI [–0.16, –0.02]). Conversely, Ts65Dn mice exhibited a significant upregulation of *Bdnf* relative to WT (WT NT vs. Ts65Dn NT, 95% CI [0.008, 0.17]). Despite these opposing patterns of dysregulation, prenatal L41 treatment effectively normalized *Bdnf* expression in both models. Following treatment, no significant differences were observed between WT and trisomic mice: in Dp1Yey mice, *Bdnf* levels in treated animals (Dp1Yey T) were comparable to WT controls (95% CI [– 0.09, 0.11]), and similarly, in Ts65Dn mice, treated animals (Ts65Dn T) showed *Bdnf* expression within the WT range (WT NT vs. Ts65Dn T, 95% CI [–0.05, 0.09]; Figure 6). These findings suggest that L41 exerts a normalizing effect on hippocampal *Bdnf* expression, irrespective of whether the initial dysregulation is an upregulation or downregulation. This highlights the potential of prenatal DYRK1A inhibition as a therapeutic approach to correct molecular alterations associated with trisomy

### Prenatal L41 Treatment Differentially Modulates NeuN- and GAD67-Positive Neuronal Populations in Dp1Yey and Ts65Dn Mice

To assess the impact of prenatal L41 treatment on neuronal development, we employed a stereological approach to quantify NeuN-positive mature neurons and GAD67-positive GABAergic interneurons in hippocampal subregions (stratum radiatum, stratum lacunosum-moleculare, and stratum moleculare). Brain sections (50 µm thick) were collected at a 1:6 sampling interval and processed for immunohistochemistry, followed by cell counting using the optical dissector method.

In both trisomic models, the number of NeuN-positive neurons was significantly elevated compared to wild-type (WT) controls (Dp1Yey NT vs. WT NT, 95% CI [821.7, 4474.8]; Ts65Dn NT vs. WT NT, 95% CI [1029.6, 3712.5]; Figure 7). Prenatal L41 treatment attenuated this increase, resulting in NeuN-positive cell counts in treated trisomic mice that were comparable to WT levels (Dp1Yey T vs. WT NT, 95% CI [–81.95, 3296.7]; Ts65Dn T vs. WT NT, 95% CI [–1227.6, 2296.8]).

**Figure 7:**
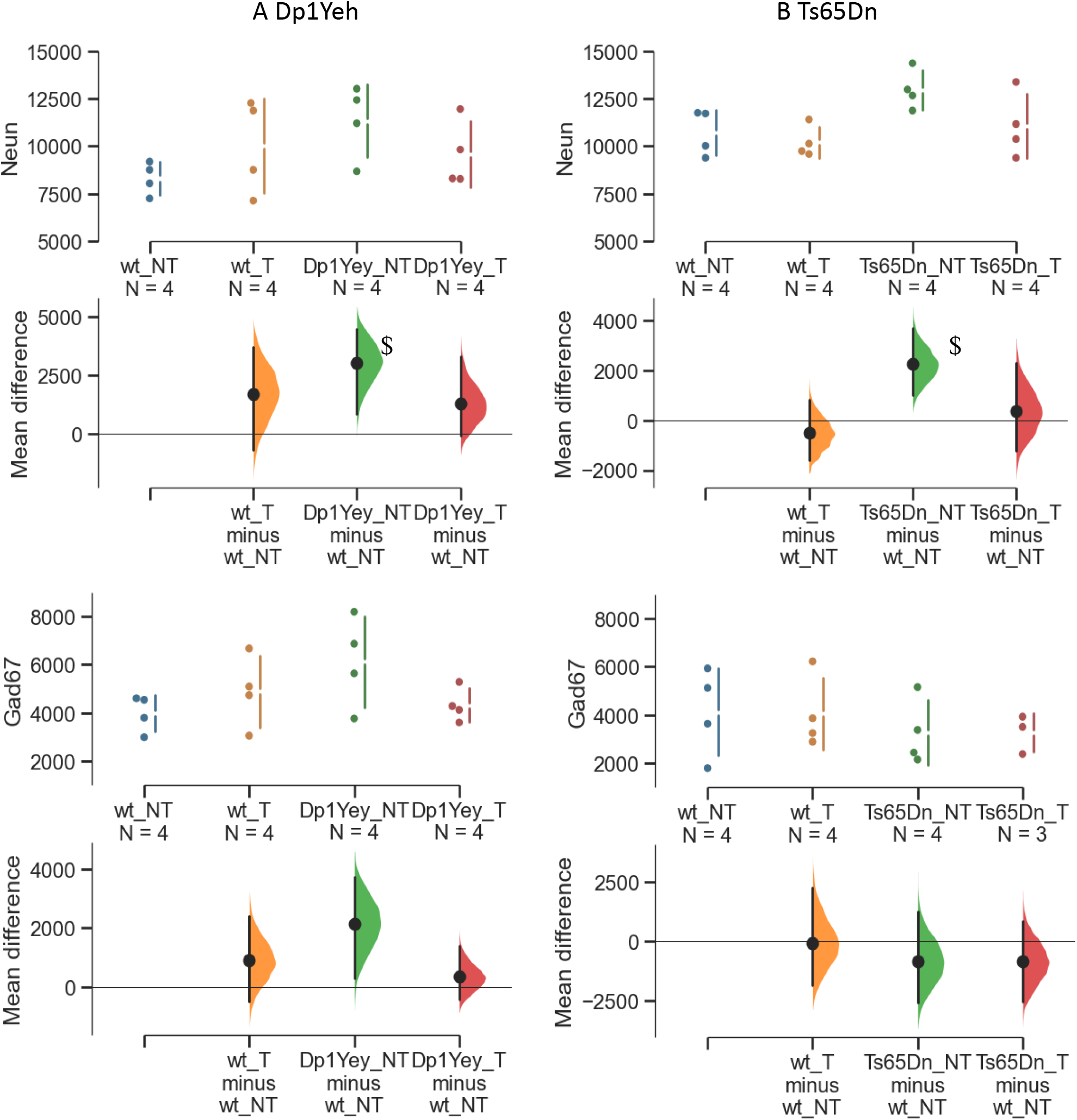
Quantification of NeuN- and GAD67-positive cells in the hippocampus of wild-type and trisomic mice, with or without prenatal L41 treatment. The prenatal L41 treatment normalizes NeuN and GAD67 expression in Dp1Yey mice, while no significant changes are observed in Ts65Dn mice. Data are presented using Gardner-Altman estimation plots. Upper panels: Raw individual cell counts are shown for each group, allowing visualization of variability and distribution. Lower panels: Bootstrap sampling distributions of the mean differences are shown for comparisons between the wild-type untreated group and each of the other groups (wild-type treated, trisomic untreated, trisomic treated). Black dots represent the observed mean differences. Horizontal black lines indicate the 95% confidence intervals, calculated from 5,000 bootstrap resamples. The symbol $ denotes that the value zero lies outside the 95% confidence interval, indicating a statistically significant difference.

For GAD67-positive interneurons, Dp1Yey mice exhibited a significant increase in cell numbers relative to WT controls (Dp1Yey NT vs. WT NT, 95% CI [299.7, 3734]). This increase was normalized following L41 treatment (Dp1Yey T vs. WT NT, 95% CI [–413, 1385]). In contrast, Ts65Dn mice showed no significant differences in GAD67-positive cell counts, regardless of treatment status.

These results demonstrate that prenatal L41 treatment selectively modulates neuronal populations in the Dp1Yey model, normalizing both NeuN- and GAD67-positive cell numbers, while its effects in Ts65Dn mice are limited. This suggests model-specific differences in responsiveness to DYRK1A inhibition during neurodevelopment.

## DISCUSSION

Here we hypothesized that in utero inhibition of DYRK1A using L41 treatment will exert long-term neurodevelopmental and cognitive benefits in two genetically distinct Down syndrome (DS) mouse models: Dp1Yey and Ts65Dn. Our findings reveal model-dependent responses to prenatal treatment, underscoring the complexity of trisomy 21 and its therapeutic modulation.

First, we confirmed that L41 crosses both the placental and embryonic blood-brain barriers, reaching the developing brain. DYRK1A kinase activity was elevated in trisomic embryos, and a single prenatal L41 injection at embryonic day 18 (E18) significantly reduced DYRK1A overactivity in Dp1Yey embryos to wild-type levels, confirming effective target engagement. However, Ts65Dn embryos exhibited high inter-individual variability, likely due to their hybrid genetic background, resulting in no consistent reduction in DYRK1A activity. Despite this, sustained kinase inhibition was observed in Dp1Yey mice from postnatal day 12 (P12) to P16, supporting L41’s potential as a prenatal therapeutic intervention.

Prenatal L41 administration differentially affected survival and behavioral outcomes in the two models. While Dp1Yey mice showed robust DYRK1A inhibition, Ts65Dn mice exhibited a potential survival benefit without significant kinase activity reduction. Importantly, the treatment with the L41 DYRK1A inhibitor was not affecting the number of viable embryos.

Behavioral assessments revealed selective cognitive improvements. Episodic memory, evaluated via the Novel Object Recognition (NOR) test, was restored in both trisomic models following L41 treatment. Working memory, assessed using the Y-maze spontaneous alternation task, was rescued in Dp1Yey mice but not in Ts65Dn mice, suggesting model-specific dependence on DYRK1A signalling. Nesting behavior and general locomotor activity remained unaltered by L41, indicating that these phenotypes may arise from DYRK1A-independent mechanisms or later developmental processes.

Ts65Dn mice displayed persistent hyperactivity across multiple tests, including the Y-maze, open field, and NOR, failing to habituate between sessions, a phenotype linked to the Mmu17-derived segment in their minichromosome (Duchon *et al*. 2022). L41 did not mitigate hyperactivity in either model, further suggesting that this trait is independent of DYRK1A overactivity.

At the molecular level, hippocampal *Bdnf* expression was dysregulated in both models but differently: downregulated in Dp1Yey, upregulated in Ts65Dn, and normalized following L41 treatment. Histological analyses revealed that NeuN-positive mature neuron counts, elevated in both trisomic models, were reduced to wild-type levels after L41 treatment. GAD67-positive GABAergic interneuron density, increased in Dp1Yey mice, was normalized by L41, consistent with prior EGCG studies. However, no significant changes were observed in Ts65Dn mice, possibly due to genetic difference in the model. These differences align with the identification of additional loci influencing behavior have been identified in DS models (Jiang *et al*. 2015; Duchon *et al*. 2021a; Duchon *et al*. 2021b; Sawa *et al*. 2022).

To ensure comparability, we employed a standardized behavioral battery, confirming that both models exhibit similar impairments in nesting, open field exploration, Y-maze performance, and NOR. However, their responses to L41 diverged, highlighting model-specific susceptibility to DYRK1A inhibition. The lack of nesting behavior improvement suggests that this phenotype may involve non-DYRK1A pathways, such as altered 5HT2A signalling (Heller et al., 2014).

Hyperactivity was more pronounced in Ts65Dn mice, as evidenced by increased arm entries in the Y-maze and greater distances travelled in OF and NOR tests. Unlike Dp1Yey mice, Ts65Dn animals failed to habituate between OF sessions and displayed persistent hyperactivity during both phases of the NOR test. These findings are consistent with previous reports linking this phenotype to the presence of the Mmu17-derived segment in the Ts65Dn minichromosome (Duchon et al., 2022; Xing et al., 2023). Importantly, L41 treatment did not mitigate hyperactivity in either model, further supporting the notion that this phenotype is independent of DYRK1A overactivity.

Our findings demonstrate that prenatal DYRK1A inhibition can partially rescue cognitive deficits in DS mouse models, with greater efficacy in Dp1Yey mice. This underscores the neurodevelopmental origins of DS-related cognitive impairments and the importance of genetic background in therapeutic responsiveness.

Limitations include small sample sizes for molecular and histological analyses and the inherent constraints of partial trisomy models. Future research should identify key DYRK1A targets during neurogenesis, particularly in GABAergic interneuron development, and clarify the temporal windows where DYRK1A inhibition is most effective. Exploring the translational potential for human DS fetuses, aiming for long-term cognitive enhancement, remains a critical goal.

Ultimately, this study provides critical insights into the neurodevelopmental mechanisms of DS and the therapeutic promise of prenatal DYRK1A inhibition, while emphasizing the need for considering model-specific approaches in therapeutic development.

## Supporting information

Statistical analysis

## Acknowledgements

We are thankful to Laurent MEIJER (Perha Pharmaceuticals) for providing Leucettine L41. We thank members of the research group, of the IGBMC laboratory and of the ICS for their help. We extend our thanks to the animal caretakers of the ICS, who are in charge of mouse wellbeing and who helped us to breed and maintain both mutant lines.

## Funding

This work was supported by Fondation Jérôme Lejeune (YH and AD), Centre National de la Recherche Scientifique (CNRS), Institut National de la Santé et de la Recherche Médicale (INSERM), the Université de Strasbourg (Unistra), and the French state funds through the Agence Nationale de la Recherche under the project PRC DYRK-DOWN ANR-18-CE16-0020 (Y.H.; J.D.G), and support from the program Investissements d’Avenir labeled IdEx Unistra (ANR-10-IDEX-0002), a SFRI-STRAT’US project (ANR 20-SFRI-0012), EUR IMCBio (ANR-17-EURE-0023) to JDG and YH, INBS PHENOMIN (ANR-10-INBS-07 PHENOMIN), the DendriDown (ANR-22-CE16-0021) and DevInDS (ANR-21-NEU2-0011) projects and the EU funded project GO-DS21 (Grant agreement ID: 848077) to YH. The funders had no role in the study design, data collection and analysis, decision to publish, or preparation of the manuscript.

## Author contribution

AD and YH conceived and designed the experiments, performed and analyzed experiments, coordinated the study and wrote the manuscript. A.D conceived and designed the experiments, performed and analyzed experiments. PG did the L41 dosage in embryo extract. JD performed the kinase activity assay. Y.H coordinated and supervised the study, and provided the financial support.

## Conflict of interest statement

AD, CC, PG, JD, and YH have declared no competing interests.

LM is the scientific director to Perha pharmaceuticals who previously developed the L41 DYRK1A inhibitor.

## Notes

### Competing Interest Statement

The authors have declared no competing interest.

